# A gut-on-chip model reveals architecture-dependent epithelial responses to microbial challenge

**DOI:** 10.64898/2026.10.08.757552

**Authors:** Manasa Nandimandalam, Puja Pandey, Mariella Mock, Mandy Holzberger, Matthias Lochner, Manuela Buettner, Andre Bleich

**Affiliations:** Institute for Laboratory Animal Science, Hannover Medical School, Hannover, Germany; Institute of Medical Microbiology and Hospital Epidemiology, Hannover Medical School, Hannover, Germany

**Keywords:** Intestinal epithelial barrier, 3D cell culture, Host–microbe interaction modeling, Gut-on-chip, Microfluidics

## Abstract

Advanced in vitro models that better recapitulate human intestinal physiology are needed to improve translational relevance and reduce reliance on animal experimentation. Microfluidic gut-on-chip systems provide controlled architectural and mechanical microenvironments that more closely resemble native tissue than conventional static cultures. However, how these features influence epithelial responses to microbial stimuli remains incompletely understood. In this study, we compared conventional two-dimensional (2D) Caco-2 monolayers with a perfused three-dimensional (3D) gut-on-chip model exhibiting crypt-like epithelial organization. Microbial challenge was employed as a biologically relevant perturbation to investigate barrier-associated responses under distinct culture conditions. Following exposure to the probiotic *Escherichia coli* Nissle 1917 (*Ec*N), epithelial morphology, barrier-associated gene expression, inflammatory signaling, and temporal response patterns were assessed over time. In a complementary approach, susceptibility to bacterial stress was examined using increasing doses of *Citrobacter rodentium* (*C. rodentium*). Both culture systems responded dynamically to microbial exposure; however, differences in response patterns were observed between static and perfused cultures. *Ec*N induced transient alterations in epithelial organization followed by recovery-associated responses, while *C. rodentium* caused dose-dependent structural and transcriptional changes. Notably, epithelial damage became apparent at lower bacterial loads in conventional 2D cultures, whereas the chip-based model maintained tissue integrity under comparable conditions. Together, these findings indicate that epithelial architecture and the surrounding culture environment influence barrier-associated responses to microbial challenge. The study highlights the utility of gut-on-chip systems for investigating host-microbe interactions and provides a framework for functional comparison of advanced intestinal in vitro models.

## 1 Introduction

The intestinal epithelium forms a highly specialized barrier between the host and the luminal environment. In addition to its central role in nutrient absorption, it regulates immune homeostasis and protects underlying tissues from microbial invasion. The intestinal barrier continuously interacts with a dense and diverse microbial community, and disturbances in this tightly regulated host-microbe relationship can contribute to the development of inflammatory and infectious diseases (1). Understanding how epithelial tissues respond to microbial stimuli is therefore essential for elucidating mechanisms of barrier maintenance, injury, and recovery.

A variety of *in vitro* models have been developed to investigate epithelial barrier function and host-microbe interactions. Among these, the human colorectal adenocarcinoma-derived Caco-2 cell line remains one of the most widely used experimental systems because of its ability to differentiate into polarized epithelial monolayers with functional tight junctions (2–4). Conventional two-dimensional (2D) Caco-2 cultures have provided important insights into epithelial permeability, barrier regulation, nutrient transport, and responses to microbial stimuli (5–7). However, static monolayer cultures do not adequately reflect several key features of the intestinal microenvironment, including three-dimensional tissue organization, extracellular matrix interactions, and luminal flow-associated mechanical forces. As a consequence, their ability to capture dynamic epithelial responses remains limited (4,5,7). To address these limitations, microfluidic organ-on-chip technologies have emerged as advanced culture platforms that more closely mimic aspects of native tissue physiology. Gut-on-chip systems combine controlled perfusion with a defined microenvironment and support the development of epithelial structures that more closely resemble the organization of the intestinal mucosa (8–10). Continuous flow further enables controlled delivery of nutrients and external stimuli while simultaneously generating mechanical conditions that are absent in conventional static cultures. Previous studies have demonstrated that such systems promote epithelial differentiation, tissue remodeling, and physiologically relevant host responses, thereby expanding the possibilities for mechanistic studies of intestinal barrier function (8,11,12).

Microbial exposure represents a biologically relevant approach for probing epithelial barrier responses under controlled experimental conditions. Different microorganisms induce distinct epithelial adaptations and stress responses, making them useful tools for investigating barrier resilience and susceptibility. *Escherichia coli* Nissle 1917 (*Ec*N) is a well-characterized probiotic strain that has been reported to support epithelial barrier integrity through modulation of tight-junction proteins and inflammatory signaling pathways (13,14). In contrast, pathogenic bacteria are capable of triggering epithelial injury and inflammatory responses. Although *Citrobacter rodentium* (*C. rodentium*) is a murine-restricted pathogen and does not naturally infect humans, it is widely used as an experimental model for studying mechanisms of epithelial stress, barrier disruption, and host responses to enteric infection (15). Under defined in vitro conditions, both microorganisms therefore provide complementary stimuli to investigate epithelial responses to microbial challenge.

In the present study, we compared conventional 2D Caco-2 monolayers with a perfused three-dimensional (3D) gut-on-chip model to investigate how culture architecture and microenvironmental conditions influence epithelial responses to microbial challenge. Following exposure to EcN, temporal changes in epithelial morphology, barrier-associated gene expression, inflammatory signaling, and TEER responses in conventional cultures were assessed. In a complementary approach, increasing doses of *C. rodentium* were used to examine susceptibility to bacterial stress in both culture systems. We hypothesized that the distinct architectural and environmental characteristics of static and microfluidic cultures would result in different epithelial response patterns following microbial perturbation. An overview of the experimental design is provided in Figure 1. Collectively, this study examines how culture format influences epithelial responses to probiotic and pathogenic bacterial challenge. By comparing conventional and microfluidic culture systems, we aim to provide further insight into the role of tissue organization and microenvironmental conditions in shaping barrier-associated responses and host-microbe interactions in advanced intestinal in vitro models.

**Figure 1:**
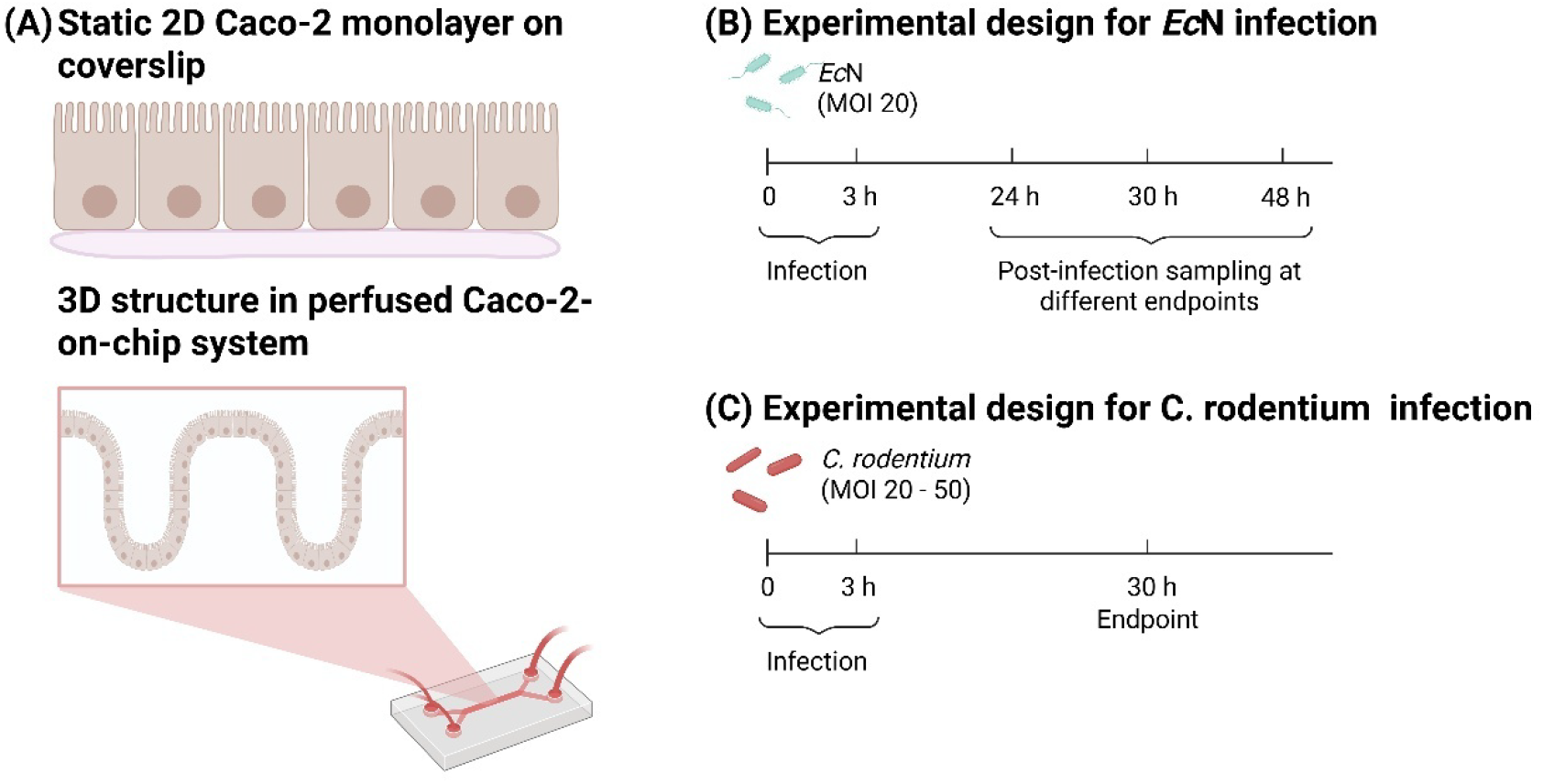
Overview of 2D and 3D Caco-2 infection models and experimental design. (A) Schematic illustration of static 2D Caco-2 monolayers cultured on coverslips and the perfused 3D microfluidic Caco-2-on-chip system supporting crypt-like epithelial architecture. (B) Experimental design for time-resolved infection with *Ec*N at MOI 20, including post-infection analysis of epithelial morphology, barrier function and gene expression under antibiotic-treated conditions. (C) Experimental design for dose-dependent infection with *C. rodentium* at MOIs 20–50, assessing epithelial damage and inflammatory responses at a defined post-infection endpoint. Experiments involving *Ec*N and *C. rodentium* were conducted independently and were not designed for direct comparison.

## 2 Materials and Methods

### 2.1 Cell culture

The human intestinal epithelial cell line Caco-2 (DSMZ, Braunschweig, Germany) was cultured in T75 flasks (Sarstedt, Germany) using Caco-2 growth medium composed of Advanced Dulbecco’s Modified Eagle Medium (DMEM) (Ref: 12634-010, Gibco, USA) supplemented with 10 % fetal calf serum (Ref: FBS.S 0615, Bio&Cell, Germany), 1 % GlutaMAX (Ref: 35050-061, Gibco, USA), and 1 % ZellShield (Ref: 13-0150, Minerva, Germany). Cells were maintained at 37 °C in a humidified incubator with 5 % CO₂ until reaching approximately 90 % confluence. For routine passaging, cells were washed with warm PBS-/- (Gibco, Germany) and detached using 0.25 % trypsin-EDTA (Ref: 0000306113, Gibco, USA) incubation for 5 minutes and sub-cultured at a 1:4 ratio every 6-7 days. The cells up to passage number 35 were considered for the experiment. For contamination control, mycoplasma test was done using Polymerase Chain Reaction (PCR) following suspicion of contamination in cell culture.

### 2.2 Static 2D culture on Coverslip

For static 2D culture, Caco-2 cells were seeded at a density of 5 × 10⁵ cells per sterile round 8 mm coverslip (Paul Marienfeld, Germany), which was placed at the bottom of each well of a 24-well plate. For quantitative PCR (qPCR) sampling, cells were seeded directly onto the well surface. Cells were maintained in Caco-2 culture medium with ZellShield as mentioned in 2.1 for 3 days prior to bacterial infection. For infection, the medium is devoid of antibiotics. Post-infection, samples were collected at 24, 30, and 48 hours for downstream analysis.

### 2.3 Static 2D culture on Transwell for TEER analysis

Caco-2 cells were seeded onto 0.4 µm pore size transwells (Corning, USA) at a density of 2 × 10⁵ cells per well and cultured for 3 days in Caco-2 medium supplemented with ZellShield. Upon reaching confluence, the medium was replaced with antibiotic-free medium, followed by bacterial infection. TEER measurements using Millicell ERS 3.0 Digital Voltohmmeter (Merk, Germany) were performed at baseline (0 hour, pre-infection), immediately after infection (3 hours), and 24 hours post-infection to assess monolayer barrier integrity.

### 2.4 3D culture with Chip system

For 3D culture, prefabricated microfluidic chips (Dynamic42, Germany) were used. Each chip contained a polyethylene terephthalate (PET) membrane with 8 µm pore size and a surface area of 1.32 cm², 200 µL of cell seeding volume. Caco-2 cells were seeded at a density of 5 × 10⁵ cells per channel into the upper chamber using Caco-2 medium with 20% FCS.

Prior to seeding, chips were sterilized with 70 % ethanol (Ref: 2212.9010, 99.0% purity, Th Geyer, Germany) for 30 minutes, then rinsed with sterile distilled water and Dulbecco’s phosphate-buffered saline containing calcium and magnesium (Ref: 14040-091, DPBS+/+, Gibco, Germany). Membranes were subsequently coated with collagen A (Ref: L7220, Lot: 0438Z, Biochrom, Germany) diluted 1:200 in DPBS+/+ for 20 minutes.

After 2 days when the cells reach confluency, the chips were connected to a perfusion system using a peristaltic pump Ismatec Reglo ICC (Ismatec, Switzerland) at a flow rate of 50 µL/min for an additional 4 days. The perfusion setup consists of connector (Chipshop, Germany) connected to one side of the chip inlet, 4 ml reservoir (Chipshop, Germany) connected to other inlet where medium is stored, and 0.5 mm diameter silicon tube (Dynamic42, Germany) connects the connector to the reservoir. When the pump starts, medium from the reservoir is perfused to the chip chamber and through silicon tube, it moves back to the reservoir forming a loop. One day prior to infection, cultures were maintained in antibiotics-free medium. On day 8, infection was performed using either *C. rodentium* or *Ec*N. Post-infection, samples were collected at 24, 30, and 48 hours for downstream analysis.

### 2.5 Experimental design

For conventional 2D culture experiments, uninfected controls and infected samples were generated in parallel within the same multiwell plate. For a given biological replicate, all conditions were cultured, infected, and processed simultaneously. Independent biological replicates were performed on separate experimental days using independently cultured cells and freshly prepared bacterial cultures.

For gut-on-chip experiments, one chip compartment served as the uninfected control, whereas the parallel compartment was exposed to the respective bacterial challenge. Control and infected conditions were therefore analyzed within the same chip experiment. Due to the duration and technical complexity of chip cultivation and perfusion, time-course experiments (24 h, 30 h, and 48 h post infection) were performed sequentially rather than simultaneously. Each time point was generated in an independent experimental run using separately prepared chips, independently cultured cells, and freshly prepared bacterial inocula. All chip experiments were conducted independently on different experimental days and each experiment was considered one biological replicate for statistical analysis.

### 2.6 Infection Model

#### 2.6.1 *Ec*N Infection

*Ec*N were cultured on Lysogeny Broth (LB) agar (Ref: X969.1, Carl Roth, Germany) supplemented with 1% Ampicillin (Ref: 0000208993, 100 mg/ml, Sigma Aldrich, USA). On the day prior to infection, *Ec*N was pre-cultured overnight in LB medium (Ref: X968.1, Carl Roth, Germany) containing 1% Ampicillin at 37°C with shaking at 180 Rotation per minute (RPM). For the infection process, pre-cultured bacteria were transferred to ice-cold LB medium and incubated for an additional 3 hours, adjusting the optical density at 600 nm (OD600) to approximately 1.0. Assuming the Caco-2 cells divide every 2.5 to 3 days, subsequently on Day 3 when the Caco-2 cells are confluent on the coverslip, the cells were infested with Log-phase *Ec*N at Multiplicity of infection (MOI) of 20 for a duration of 3 hours and incubated at 37°C. Following the infection period, the medium was replaced with fresh medium with antibiotics to eliminate non-adherent bacteria and incubated overnight at 37°C for up to 24-, 30- and 48-hours post-Infection. In the chip system, *Ec*N infection was done through reservoir where the bacteria are passed on to the cells in the chip chamber during perfusion. The cover slips were fixed with ice-cold methanol for 10 minutes and used for immunohistochemistry (IHC) staining. The infected Caco-2 cells in the wells were isolated using RNA-free lysis buffer for qPCR.

The chip was removed from the perfusion system, and the membrane surface was prepared for IHC staining and quantitative PCR (qPCR) (Thermofischer, USA) analysis to evaluate infection outcomes and cellular responses.

#### 2.6.2 *C. rodentium* infection

In order to analyze the effect of pathogenic bacteria, *C. rodentium* was introduced to our 2D and 3D Caco-2 system. *C. rodentium* was precultured on 5 ml LB medium overnight at 37°C, 180 RPM. For the infection, *C. rodentium* was further cultured on an additional 50 ml LB medium till OD600 was around 1. The cells were infected with *C. rodentium* MOI 20, 30, 40, and 50 in the absence of antibiotics. The infection was done for 3 hours and changed to medium with ZellShield, which was further cultured for 30 hours. The cells were used for staining or for qPCR.

### 2.7 Immunocytochemistry

For the structural characterization, the Caco-2 cells were stained for tight junction. The cells on both chip and coverslip were fixed in ice cold methanol (Ref: 1462.1011, Batch: 30.1981006, 99.7% purity, Th Geyer, Germany) for 10 minutes. The membrane of the chip was carefully cut out from the chip and transferred to 24-well plates containing PBS as the membrane should not get dry. Similarly, the further steps with coverslips were done on paraffin paper. Blocking and permeabilization was done using 10 % goat serum (Ref: 16210064, Gibco, USA) diluted in PBS containing 0.1 % TritonX (Ref: S26-36-23, Sigma-Aldrich, USA) for 30 minutes. Incubation with 1:50 rabbit anti-ZO-1 polyclonal antibody (catalogue number: 61-7300, Lot: XL361222, 0.25 mg/ml Invitrogen, USA) diluted in 100 µl blocking and permeabilization buffer was done overnight at 4°C. Following the washing step, Alexa Fluor 555 anti-rabbit, polyclonal, (catalogue number: 406412, Lot: B383581, 0.5 mg/ml, Biolegend, USA) was applied to detect the primary antibody binding. Nuclei were counterstained with VECTASHIELD antifade mounting medium with DAPI (Ref: H1200 and Lot: ZK1102, Vector Laboratories, USA). The imaging was done using fluorescence microscope Axio Imager.M2 (Zeiss, Germany) with AxioCam MR R3 camera. Image capture and editing was done using Zen 3.9 Zen Lite software. Images were captured using lenses 10x with 0.3 numerical aperture, and 20x with 0.5 numerical aperture from Zeiss. Fluorescent imaging was performed using Alexa fluor and Fluorescein dyes under appropriate excitation and fluorescent.

### 2.8 qPCR

For qPCR, the cells were harvested, lysed in RNA Quick-RNA™ Micro Prep Kit Lysis Buffer (Zymo Research, USA) and stored at -20°C for further use in RNA isolation. RNA isolation was done using RNA Quick-RNA™ Micro Prep Kit (Ref: E1009-A, Zymo Research, USA) as per manufacturer’s instructions. For cDNA synthesis, RNA was reverse transcribed using the QuantiTect® Reverse Transcription Kit (Ref: 205313, Qiagen, Germany) according to the manufacturer’s protocol. QuantStudio™ 6 Flex Real-Time PCR System (Applied Biosystems™, USA) was used for qPCR. TaqMan®-based assay primers (ACTB: Hs01060665_g1; TNF: Hs00174128_m1, OCLN: Hs05465837_g1, MKI67: Hs04260396_g1, TJP1: Hs01551871_m1 (Thermo Fischer Scienctific, USA) were used. ACTB was used as a housekeeping gene. Analysis was done in triplets. Relative gene expression was calculated using the 2−ΔΔCt method. Expression values were normalized to ACTB and are presented relative to the corresponding reference sample, which were set to 1.

### 2.9 Statistical Analysis

GraphPad Prism8® software (San Diego, USA) is used for the statistical analysis. The statistical significance is carried out using One way-Anova, Sidak test. The outliers were identified with Rout method with false discovery rate (Q) of 1%. P value ≤0.05 is considered significant. The data are expressed as standard error of mean (SEM). The number of independent biological replicates for each experiment is indicated in the corresponding figure legends.

## 3 Results

### 3.1 Time-resolved epithelial responses to probiotic *Ec*N in 2D and 3D Caco-2 systems

To investigate temporal effects of *Ec*N on epithelial barrier-associated responses, Caco-2 monolayers and chip-cultured epithelia were exposed to *Ec*N (MOI 20) for 3 h and analyzed over a period of 48 h. In 2D monolayers, immunofluorescence staining of the tight-junction protein ZO-1 revealed alterations in epithelial organization between 24 and 30 h post infection, characterized by the appearance of intercellular gaps and partial disruption of junctional continuity. At 48 h post infection, ZO-1 staining appeared more continuous, indicating partial restoration of epithelial organization (Fig. 2A). TEER measurements supported these observations. Baseline TEER values were comparable between groups and remained largely unchanged immediately after infection. At 24 h post infection, *Ec*N-treated monolayers showed a significant reduction in TEER compared with uninfected controls (Fig. 2B). Gene expression analyses demonstrated temporal variation in barrier- and inflammation-associated markers. *KI67* expression remained largely unchanged throughout the observation period. *OCLN* and *TJP1* expression showed variation over time, with numerically higher expression levels at later time points, whereas *TNF* expression displayed a transient increase during the infection course (Fig. 2C).

**Figure 2:**
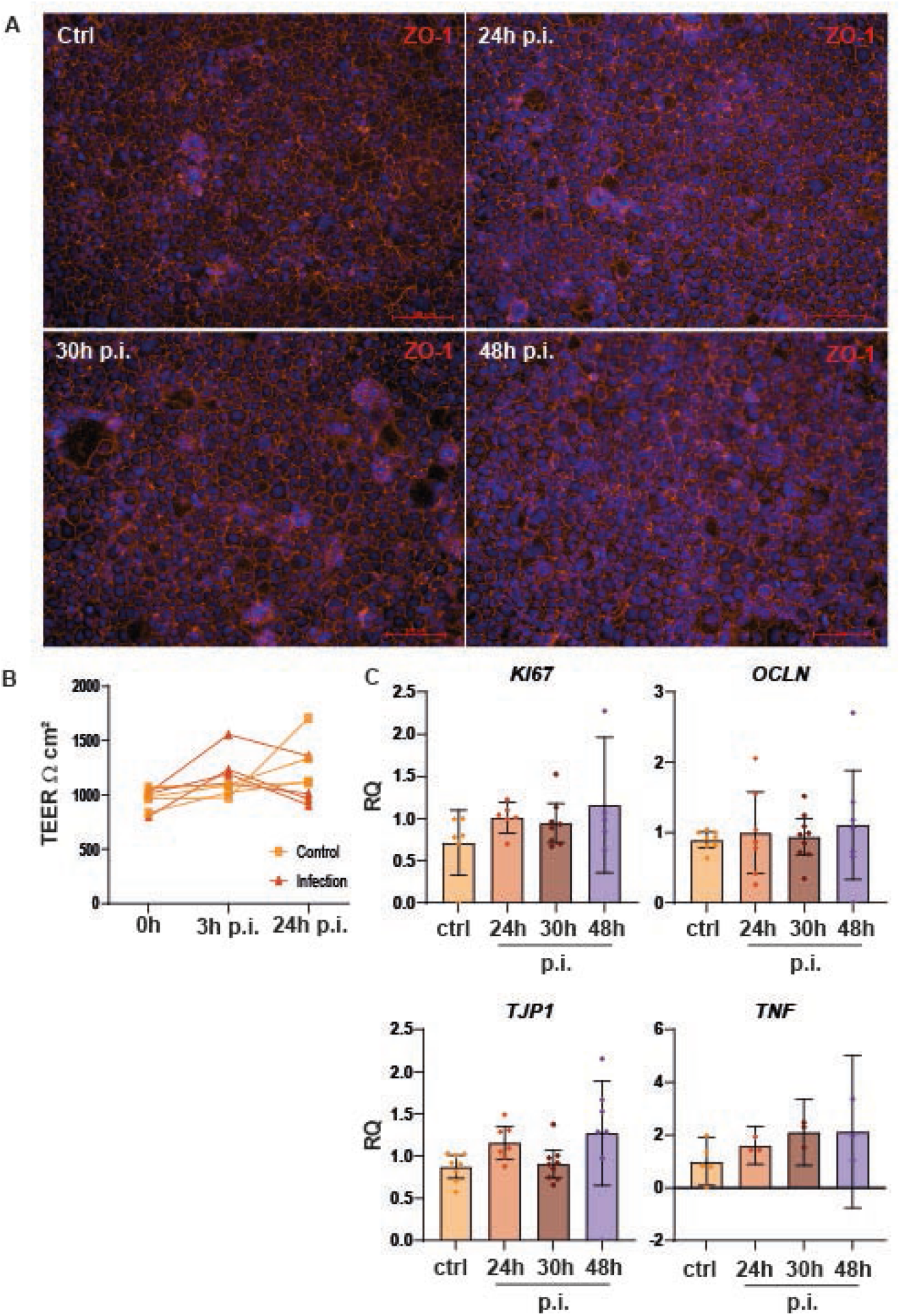
Temporal epithelial responses to *Ec*N in conventional 2D Caco-2 cultures. (A) Immunofluorescence staining of the tight junction protein ZO-1 in uninfected control monolayers and at 24, 30 and 48 hours following *Ec*N infection (MOI 20), showing transient junctional disruption and subsequent restoration of epithelial continuity. (B) Transepithelial electrical resistance (TEER) measurements over time demonstrating a reduction in barrier function at 24 hours post-infection, followed by recovery. Data represent four independent experiments. (C) qPCR analysis of *KI67*, *OCLN*, *TJP1*, and *TNF* expression during the course of *Ec*N infection. Expression levels are shown relative to uninfected controls. Data represent up to nine independent biological experiments (mean ± SEM). Statistical analysis was performed using one-way ANOVA; no significant differences were observed.

In the perfused 3D gut-on-chip model, *Ec*N exposure was associated with temporal changes in epithelial morphology and gene expression. At early time points, crypt-like structures remained visible and *OCLN* and *KI67* expression showed only minor variation. At later time points, localized alterations in epithelial organization were observed together with changes in *TJP1* and *TNF* expression, while epithelial structures remained detectable throughout the observation period (Fig. 3).

**Figure 3:**
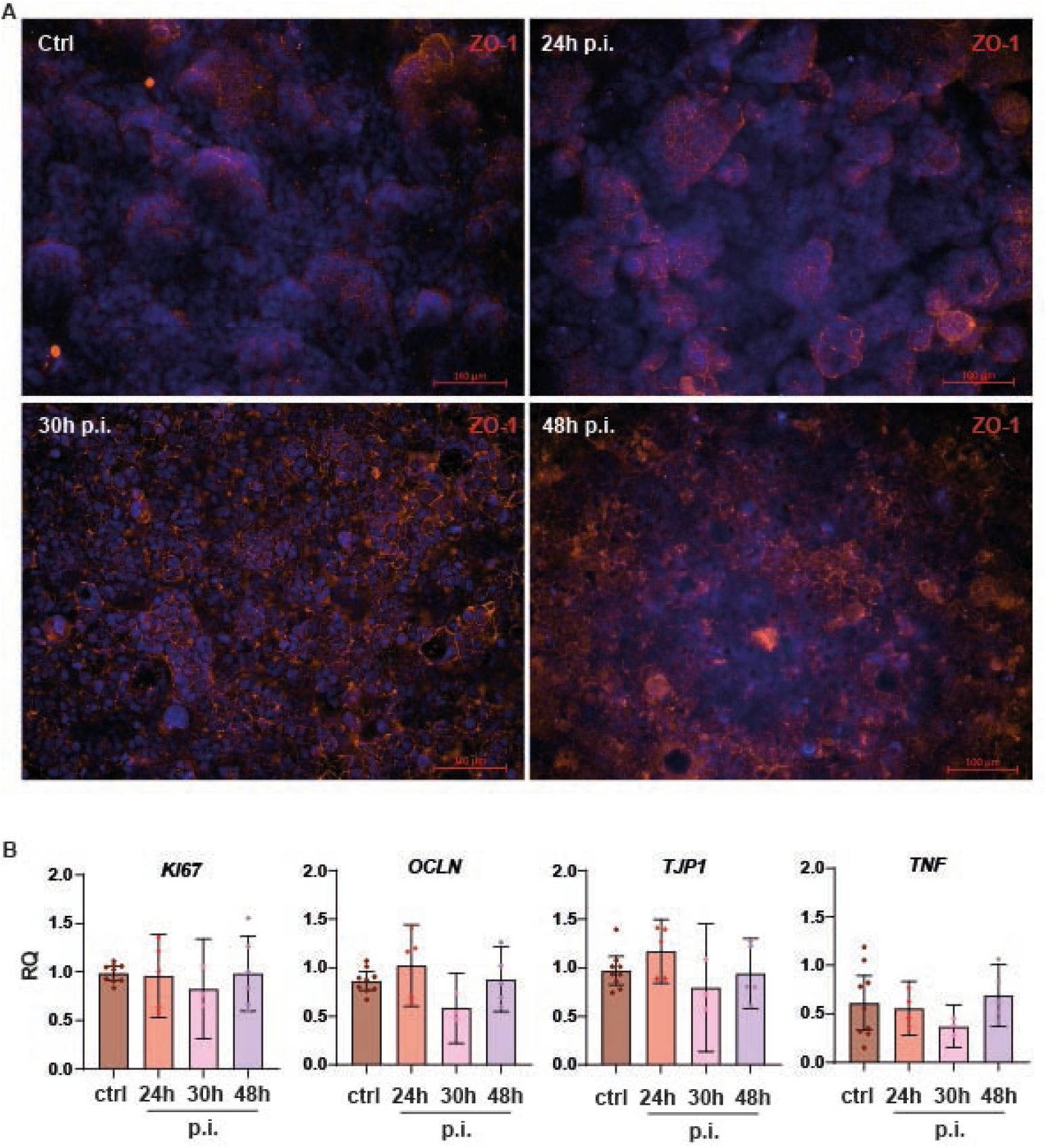
Time-dependent epithelial responses during *Ec*N infection in the 3D Caco-2 gut-on-chip model. (A) Representative immunostaining images of chip cultured epithelia at 24, 30 and 48 hours post *Ec*N infection (MOI 20), illustrating preserved crypt like structures at early time points and transient epithelial injury followed by regeneration. (B) qPCR analysis of *OCLN*, *TJP1*, *KI67*, and *TNF* expression at 24, 30, and 48 h following *Ec*N exposure in the gut-on-chip model. Expression levels varied over the observation period but did not reach statistical significance. Relative gene expression was normalized to uninfected controls. Data are shown as mean ± SEM from up to nine independent biological experiments. Statistical analysis was performed using one-way ANOVA.

### 3.2 Architecture-dependent epithelial damage induced by pathogenic *C. rodentium*

To assess responses to pathogenic bacterial challenge, both culture systems were exposed to increasing doses of *C. rodentium* (MOI 20-50). In conventional 2D monolayers, no obvious structural alterations were observed at MOI 20. Beginning at MOI 30, increasing disruption of epithelial organization became apparent, with widening intercellular gaps and progressive loss of junctional continuity at higher bacterial loads (Fig. 4A).

**Figure 4:**
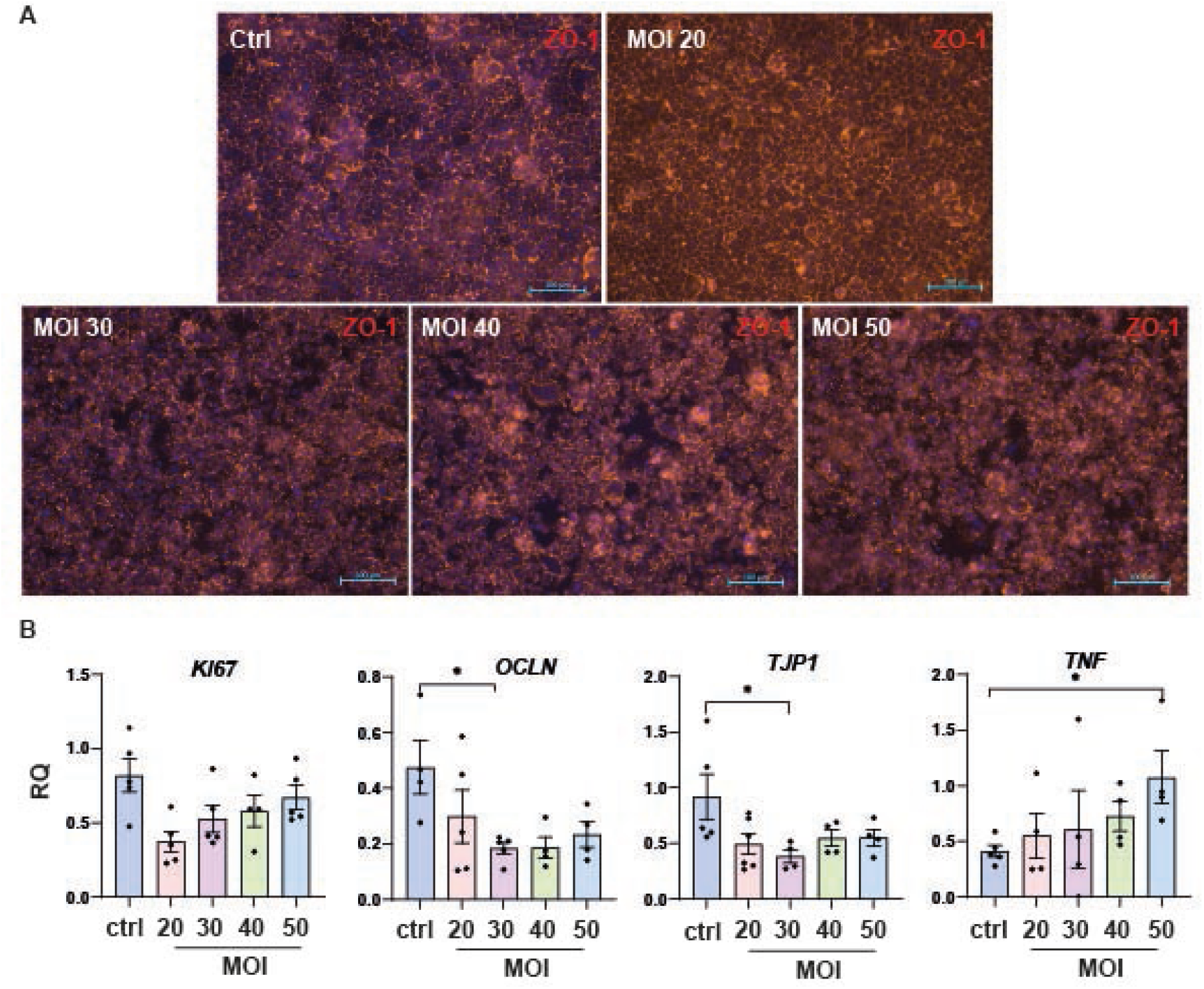
Dose dependent disruption of epithelial integrity by *C. rodentium* in 2D Caco 2 monolayers. (A) Immunofluorescence images of tight junction staining in monolayers infected with *C. rodentium* at MOIs 20, 30, 40 and 50, showing progressive loss of junctional continuity with increasing bacterial load. (B) qPCR analysis of *TJP1*, *OCLN*, *KI67*, and *TNF* expression in response to increasing *C. rodentium* loads (MOI 20-50) in conventional 2D Caco-2 cultures. Relative expression values were normalized to uninfected controls. Data are presented as mean ± SEM from up to five independent biological experiments. Statistical analysis was performed using one-way ANOVA; statistically significant differences are indicated where applicable.

In the gut-on-chip model, crypt-like epithelial structures remained preserved up to MOI 30. Detectable structural alterations emerged at MOI 40 and were more extensive at MOI 50, including areas of epithelial detachment and loss of structural continuity (Fig. 5A). Gene expression analyses revealed changes in barrier-associated and inflammatory markers following *C. rodentium* exposure. *TJP1* and *OCLN* expression varied across infection conditions, whereas *KI67* expression remained relatively stable. *TNF* expression increased following bacterial challenge in both culture systems, with the highest values observed at elevated bacterial loads (Figs. 4B and 5B).

**Figure 5:**
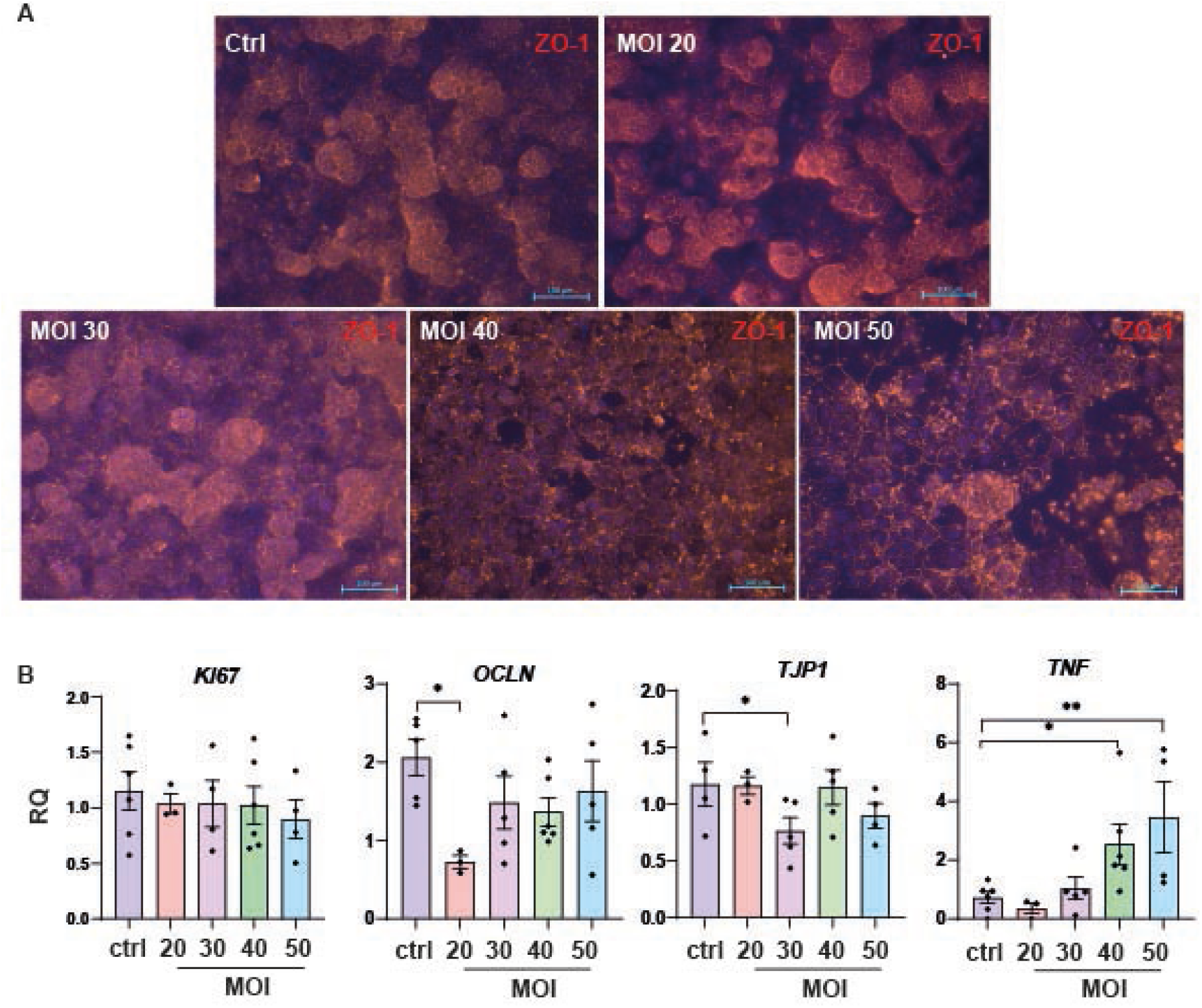
Epithelial responses to increasing *C. rodentium* challenge in the perfused 3D Caco-2 gut-on-chip model. (A) Representative images of chip cultured epithelia infected with *C. rodentium* at MOIs 20, 30, 40 and 50, showing preserved crypt like structures up to MOI 30 and detectable epithelial damage emerging at MOI 40. (B) qPCR analysis of *TJP1*, *OCLN*, *KI67*, and *TNF* expression following exposure to increasing doses of *C. rodentium* (MOI 20-50) in the gut-on-chip model. Relative gene expression was normalized to uninfected controls. Data are shown as mean ± SEM from six independent biological experiments. Statistical analysis was performed using one-way ANOVA. Statistically significant differences are indicated in the figure.

## 4 Discussion

Microfluidic gut-on-chip systems are increasingly used to study intestinal epithelial biology under conditions that more closely resemble aspects of the native intestinal microenvironment than conventional static cell cultures. In the present study, we compared epithelial responses to microbial challenge in conventional 2D Caco-2 monolayers and a perfused 3D gut-on-chip model in order to examine how differences in culture architecture and microenvironmental conditions influence barrier-associated responses. Exposure to *Escherichia coli* Nissle 1917 (*Ec*N) resulted in temporal changes in epithelial morphology and gene expression in both culture formats. In 2D cultures, alterations in ZO-1 organization were accompanied by a transient decrease in TEER, while later time points showed a more continuous junctional staining pattern. Changes in the expression of tight-junction-associated and inflammatory genes were observed over the course of the experiment, whereas *KI67* expression remained relatively stable. Together, these findings suggest that epithelial adaptation to *Ec*N exposure occurred primarily through modulation of barrier-associated pathways rather than through marked changes in proliferative activity. These observations are consistent with previous reports describing effects of *Ec*N on epithelial barrier regulation and inflammatory signaling (16,17). In the gut-on-chip model, temporal alterations in epithelial organization and gene expression were likewise observed following *Ec*N exposure. The three-dimensional epithelial structures remained detectable throughout the observation period, although localized morphological changes were evident at later time points. Compared with conventional monolayers, the chip system enabled the assessment of epithelial responses within a more complex architectural context that incorporates extracellular matrix support and continuous perfusion. While the present data do not allow conclusions regarding the contribution of individual factors, they suggest that culture environment can influence the manifestation of epithelial responses to microbial stimulation.

Differences between the two systems became particularly apparent following exposure to *Citrobacter rodentium*. In static monolayers, morphological alterations became visible at lower bacterial loads, whereas chip-cultured epithelia maintained overall structural organization up to higher infection doses. These observations indicate that epithelial responses to microbial stress differ between the investigated culture formats. However, because the systems differed not only in tissue architecture but also in membrane properties, extracellular matrix coating, serum concentration, culture duration, and perfusion conditions, the specific contribution of any individual parameter cannot be determined from the present study. Future studies employing systematically matched culture conditions will be necessary to dissect the relative impact of these variables. Several additional limitations should be considered when interpreting the findings. First, *C. rodentium* is a murine-adapted pathogen and therefore does not fully reflect human-specific host-pathogen interactions. The current study was not intended to model human infection but rather to investigate epithelial responses to defined microbial perturbations. Second, functional barrier measurements were only available for conventional 2D cultures, whereas assessment of the gut-on-chip model relied primarily on morphological and transcriptional endpoints. The inclusion of permeability assays or chip-based TEER measurements would provide additional information regarding barrier function and should be addressed in future studies. Third, the use of an immortalized epithelial cell line does not fully capture the cellular complexity of the intestinal mucosa, including immune, stromal, and microbial community interactions. Despite these limitations, the study demonstrates that microbial exposure can serve as a useful experimental stimulus for investigating epithelial responses in different intestinal culture systems. The combination of morphological observations, barrier-associated measurements, and transcriptional analyses revealed distinct response patterns between static and microfluidic cultures and highlights the importance of considering culture context when interpreting host-microbe interactions *in vitro*.

In conclusion, our findings indicate that epithelial responses to microbial challenge are influenced by the structural and environmental characteristics of the culture system. The observed differences between conventional monolayers and a perfused gut-on-chip model underscore the importance of tissue organization and microenvironmental conditions in shaping barrier-associated responses. These results contribute to the ongoing development of advanced intestinal *in vitro* models and support their application in studies of epithelial barrier biology and host-microbe interactions.

## 6 Author Contributions

M.B., M.N. and P.P. conceived and designed the experiments and wrote the manuscript. M.B., M.N. and P.P. and M.H. designed pictures. M.N. and P.P. and M.M. performed the experiments and analyzed the data. A.B. and M.L. provided access to materials and infrastructure. M.B. supervised the work.

## 8 Acknowledgments

No competing interest was reported by the authors(s).

## 9 Generative Artificial Intelligence (AI)

The use of Microsoft Copilot (GPT-5) was fully disclosed in the manuscript. The tool was used exclusively for language editing and improvement of readability. All scientific content, analyses, interpretations, and conclusions were developed, verified, and approved by the authors.

## 10 Data Availability Statement

The raw data supporting the conclusions of this manuscript are available from the corresponding author on reasonable request to any qualified researcher.

